# Antimicrobial silver inhibits bacterial movement and stalls flagellar motor

**DOI:** 10.1101/2020.02.19.956201

**Authors:** Benjamin Russell, Ariel Rogers, Matthew Kurilich, Venkata Rao Krishnamurthi, Jingyi Chen, Yong Wang

## Abstract

Silver (Ag) has been gaining broad attention due to their antimicrobial activities and the increasing resistance of bacteria to commonly prescribed antibiotics. However, various aspects of the antimicrobial mechanism of Ag have not been understood, including how silver affects the motility of bacteria, a factor that is intimately related to bacterial virulence. Here we report our study on the antibiotic effects of Ag^+^ ions on the motility of *E. coli* bacteria using swimming and tethering assays. We observed that the bacteria slowed down dramatically when subjected to Ag^+^ ions, providing direct evidence showing that Ag inhibits the motility of bacteria. In addition, through tethering assays, we monitored the rotation of flagellar motors and observed that the tumbling frequency of bacteria increased significantly in the presence of Ag^+^ ions. Furthermore, the rotation of bacteria in the tethering assays were analyzed using hidden Markov model (HMM); and we found that Ag^+^-treatment led to a significant decrease in the tumbling-to-running transition rate of the bacteria, suggesting that the rotation of bacterial flagellar motors was stalled by Ag^+^ ions. This work provided a new quantitative understanding on the mechanism of Ag-based antimicrobial agents in bacterial motility.

## Introduction

The rising prevalence of antibiotic-resistance in harmful microbes due to overuse of conventional antibiotics has become a serious global concern for public health[1, 2, 3], posing the need for different approaches for fighting against drug-resistant microbes[4, 5]. Recent research in the past two decades revisited the antimicrobial activities of noble metals, such as silver (Ag), in different forms – including ions and nanoparticles – and has uncovered their strong capacity for suppressing bacterial growth and killing bacteria[6, 7, 8]. Exciting progress has been made towards understanding the antimicrobial mechanism of Ag, suggesting that Ag caused multidirectional damages to bacteria, including DNA damage, membrane disruption, free radical generation (ROS), and loss of ATP production[7, 9, 10, 11, 12, 13]. However, various aspects of the antimicrobial mechanism of Ag remain elusive, especially that the temporal resolution for understanding the Ag-caused damages in bacteria Ag is still limited[7, 14, 15]. This includes how silver affects the motility of bacteria, which is tightly coupled to bacterial virulence[16].

Motility is essential to many bacteria for detecting and pursuing nutrients, as well as avoiding and fleeing from toxicants. Certain bacteria, such as *Escherichia coli* (*E. coli*), use flagella to move in aqueous environments[17]. *E. coli* flagella are filaments extending outward from the bacteria[18]. The flagella are connected to and driven by motors embedded in the bacterial membrane through hooks[19]. For *E. coli* – peritrichous bacteria with flagella covering their entire surfaces, their movement depends on the rotation direction of their flagella[17, 19]. When flagella rotate counterclockwise (CCW), they are bundled and propel the bacteria to move directionally (i.e., running) for purposeful movement toward chemical attractants or away from repellents[17, 20]; when flagella rotate clockwise (CW), they are splayed out, resulting in reorientation (i.e., tumbling) of the bacteria[17, 20]. The *E. coli* flagella contains mainly three parts: the filament, the hook, and the basal body[17, 20]. The basal body consists of several rings, some of which (e.g., MS ring and C ring) are essential components of the flagellar motor for driving the rotation of the flagella[17]. Structurally, the flagellar motor involves both the stator proteins (e.g., MotA and MotB) and the rotor proteins (e.g., FliG, FliM, and FliN), which also play critical roles in the torque generation of the motor[17]. Functionally, the CW/CCW direction of the flagellar motor’s rotations relies on another set of chemotaxis proteins (e.g., CheY, CheZ, CheA, CheW, CheR, and CheB). For example, the flagellar motor switches from CCW rotation to CW rotation when the phosphorylated response regulator CheY binds to the flagellar motor[21].

As Ag in various forms (e.g., ions, nanoparticles) suppresses and kills bacteria, we hypothesized that the motility of bacteria is significantly affected by Ag. This hypothesis is indirectly supported by evidence from previous studies. For example, Ivask et al. performed liquid-culture-based high-throughput growth assays for a library of single-gene-deletion strains of *E. coli*, and found that a series of flagella-related mutants (e.g., *fliG*, *fliM*, *flgF*, *flgG*, etc., which are involved in the assembly and function of flagella) were sensitive to Ag^+^ ions and Ag nanoparticles[13]. Also, plate-based chemical genetic screening assays on a similar library identified and confirmed some flagella-related genes (e.g., *flgA*, *flgD*, *flgJ*, *flgK, fliC*, *fliE*, *fliL*, *fliP*, *fliR*, and *motB*)[22]. In addition, recent work by us and others showed that Ag affects the organization and function of certain universal regulatory proteins in bacteria, such as histone-like nucleoid structuring (H-NS) proteins, which regulate bacterial chemotaxis and motility[14, 22, 23, 24, 25]. Furthermore, although mixed results were present, plate-based swimming and swarming motility assays suggested that Ag could change the motility of bacteria under certain conditions[26]. On the other hand, Ag^+^ ions have been used for staining bacterial flagella for decades[27], implying that Ag^+^ ions interact with flagella.

However, few studies on real-time observation and quantification of Ag’s effects on bacterial movement are presented in the literature[7, 28]. In this work, we investigated the antibiotic effects of Ag^+^ ions on the swimming behavior of *E. coli* bacteria based on microscopic imaging, with a temporal resolution of 15–50 ms. Ag^+^ ions were chosen for two reasons. First, Ag^+^ ions are effective at suppressing and killing bacteria[6, 10, 29]. Second, the release of Ag^+^ ions from AgNPs is one major contribution to the toxicity of AgNPs[7]. Through the swimming assays, we provided direct evidence showing that Ag inhibits the motility of bacteria. In addition, we monitored the rotation of flagellar motors of E. coli bacteria though tethering assays in the absence and presence of Ag^+^ ions, directly observing that Ag^+^ ions increased the frequency of bacterial tumbling. Furthermore, based on hidden Markov model (HMM) analysis, we found that Ag^+^-treatment caused bacterial transition rate from the tumbling state to the running state to decrease significantly, suggesting that the rotation of bacterial flagellar motors was stalled by Ag^+^ ions. This real-time quantification analysis by high temporal resolution microscopic imaging provides direct evidences of Ag effects on bacterial mobility.

## Materials and Methods

### Bacterial strain and growth

An *E. coli* K12-derived strain from Refs.[15, 23, 30, 31] was used in this study. The strain has been used in previous investigations of the antimicrobial activities of Ag^+^ ions and AgNPs[15, 23, 31]. This strain has the *hns* gene knocked out from the chromosomal DNA, but supplemented with a plasmid encoding for the H-NS protein fused to mEos3.2 fluorescent protein[32] and for resistance to kanamycin and chloramphenicol[15, 23, 30, 31].

Each experiment started with inoculating a single bacterial colony into 5 mL of Luria Broth (LB) medium supplemented with kanamycin and chloramphenicol (50 µg/mL and 34 µg/mL, respectively)[23]. The liquid culture was grown at 37°C in a shaking incubator (250 RPM) overnight. On the second day, the overnight culture was diluted by 5000× into 5mL of fresh LB medium with the antibiotics. The new culture was grown at 32°C[33, 34, 35] in the shaking incubator until the bacterial culture reached the mid-exponential phase (OD_600_ ≈ 0.3), followed by measurements as described below.

### Phase contrast microscopy

Measurements in the swimming and tethering assays were done at room temperature using phase contrast microscopy on an Olympus IX-73 inverted microscope equipped with a 100×, NA=1.25 phase-contrast, oil-immersion objective (Olympus) and an EMCCD camera (Andor Technology). The microscope and data acquisition was controlled using Micro-Manager[36, 37]. The effective pixel size of recorded images/movies was 0.16 µm.

### Swimming assay

In swimming assay experiments, *E. coli* bacteria at OD_600_ ≈ 0.3 were treated with Ag^+^ ions at 30 µM or 40 µM for 1, 2, and 4 hr, which clearly showed suppressed growth. At each time point, 2 mL of the bacterial culture were transferred to a cleaned glass-bottom Petri-dish, followed by monitoring and recording the free swimming of the bacteria using phase-contrast microscopy. The swimming of untreated bacteria (i.e., before the addition of Ag^+^ ions, or 0 hr) was monitored and used as negative controls. The exposure time was set to 30 ms, while the actual time interval between adjacent frames of the acquired movies was 54 ms. The acquired movies of freely swimming bacteria were processed in ImageJ by inversion, smoothing, and background subtraction[38, 39], followed by automated identification and localization of the bacteria using custom-written MATLAB programs[40]. The localizations of the bacteria were then linked into trajectories following standard algorithms[40, 41, 42], using a maximum displacement between adjacent frames of 1.92 µm (12 pixels), a memory of 0 frame (i.e., no gap), and a minimum length of 12 frames. The identified trajectories further went through a manual quality control process by removing the bacteria that were stuck on the glass surface or formed large clumps.

The trajectories of the bacteria in the freely swimming assays were further analyzed using custom-written or open-source Python programs. For example, the instantaneous velocities were calculated from the trajectories **r**(*t*) of the bacteria, 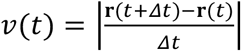, where *Δt* = 54 ms. In addition, we estimated the maximum chord-to-arc ratio 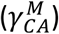 for each trajectory, inspired by TumbleScore[43], 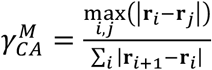, where **r**_*i*_ and **r**_*j*_ were positions of a single trajectory. Furthermore, the changing rates of swimming directions *Ω* were estimated directly from the trajectories[43, 44, 45], 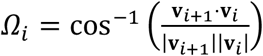. Lastly, we calculated the ensemble mean-square-displacement (MSD) for each sample using the *trackpy* Python package[42], MSD(*τ*) = ⟨(**r**(*t* + *τ*) − **r**(*t*))^2^⟩, where *τ* is the lag time.

### Tethering assay

In tethering assay experiments[46, 47], *E. coli* bacteria in the mid-exponential phase (OD_600_ ≈ 0.3) were tethered to glass-bottom Petri-dishes through their flagella. The tethering was achieved by coating the glass surface with biotinylated BSA, neutravidin, and biotinylated anti-FliC antibody sequentially[48, 49]. *E. coli* flagella bind to the anti-FliC antibody[50, 51], immobilizing the bacteria. The rotations of the tethered bacteria were monitored and recorded under phase contrast microscopy with an exposure time of 5 ms for 10000 frames without Ag^+^ ions (the actual time interval between adjacent frames was 14.1 ms). Then Ag^+^ ions were directly added to the Petri-dish at a final concentration of 40 µM, followed by recording the rotations of the same bacteria for 50,000 to 100,000 frames. For negative controls, LB medium (instead of Ag^+^ ions) was added to the Petri-dish and the rotations of the bacteria were recorded similarly. More than 10 replicate experiments were performed independently on different days.

Bacteria in the tethering assays were identified and characterized in each frame of the recorded movies using custom-written Python programs based on the *scikit-image* package[52]. From the primary axis of the identified bacteria, the orientation *θ* of the bacteria were obtained[53, 54], followed by estimating the angular velocities of the bacterial rotation *ω* = *Δθ*/*Δt*, where *Δt* = 0.0141 s. Note that frames containing other non-tethered bacteria invading the region of the tethered ones were removed from further analysis to ensure accuracy. The *ω*-trajectories were analyzed using the hidden Markov model (HMM)[55], in which two states of the bacteria (running and tumbling) were assumed. In addition, Gaussian emission distributions were applied for the emission from the two states to the observable (i.e., angular velocities *ω*)[55]. The HMM analysis was done using the *hmmlearn* Python package. For each bacterium in the tethering assay, we fitted the HMM model using the *ω*-trajectory before the addition of Ag^+^ ions (or LB medium). Then the fitted model was used to predict the states for the data after the addition of Ag^+^ ions (or LB medium), from which the probabilities of the two states and the transition rates were estimated[56].

## Results

### Lower motibility of bacteria caused by Ag^+^ ions

We first examined the effects of Ag^+^ ions on the motility of *E. coli* bacteria using swimming assays[57, 58, 59]. When *E. coli* bacteria at OD_600_ ≈ 0.3 were treated with Ag^+^ ions at 40 µM for 1, 2, and 4 hr, the cell density did not increase and the bacterial growth was suppressed. At each time point, 2 mL of the bacterial culture were taken to a glass-bottom Petri-dish, followed by monitoring and recording the free swimming of the bacteria using phase-contrast microscopy. Untreated bacteria (i.e., 0 hr) were measured as negative controls, and we observed that the treated bacteria were much slower (Movies M1 and M2). From the movies of the freely swimming bacteria, the trajectories **r**(*t*) of individual bacteria were obtained. 200 randomly chosen examples of trajectories for each experimental condition were shown in Fig. 1A, where longer traveling distances were observed for the untreated bacteria compared to the ones treated with Ag^+^ ions. To see this difference more clearly, we plotted the corresponding rose graphs[43], in which the displacements of the bacteria from their individual initial positions were drawn, *Δ***r**^0^(*t*) = **r**(*t*) − **r**(0). 300 randomly chosen examples were shown in Fig. 1B, where the first 12 frames of the trajectories were shown to eliminate the differences due to different lengths of trajectories[43]. It is obvious that the motility of bacteria decreased significantly after the treatment with Ag^+^ ions. Note that, although the trajectories are longer (up to ~70 frames), only the first 12 frames were used in the rose graphs (Fig. 1B) to make direct comparisons. We quantified the mean and 90^th^ percentile of the displacements of the first 12 frames of all the trajectories in each condition, shown as solid and dotted circles in the rose graphs (Fig. 1B), respectively. We found that the two circles for untreated bacteria did not change significantly from 0 to 4 hr, indicating that the motility of the bacteria remained similar. In contrast, the treated bacteria showed much smaller radii for both the mean 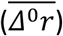 and 90^th^ percentile circles, indicating Ag^+^-treatment led to lower bacterial motility. We also note that the radii slightly increased for longer treatment time, implying possible recovery of the bacteria as reported by our previous results[6, 15].

**Figure 1.**
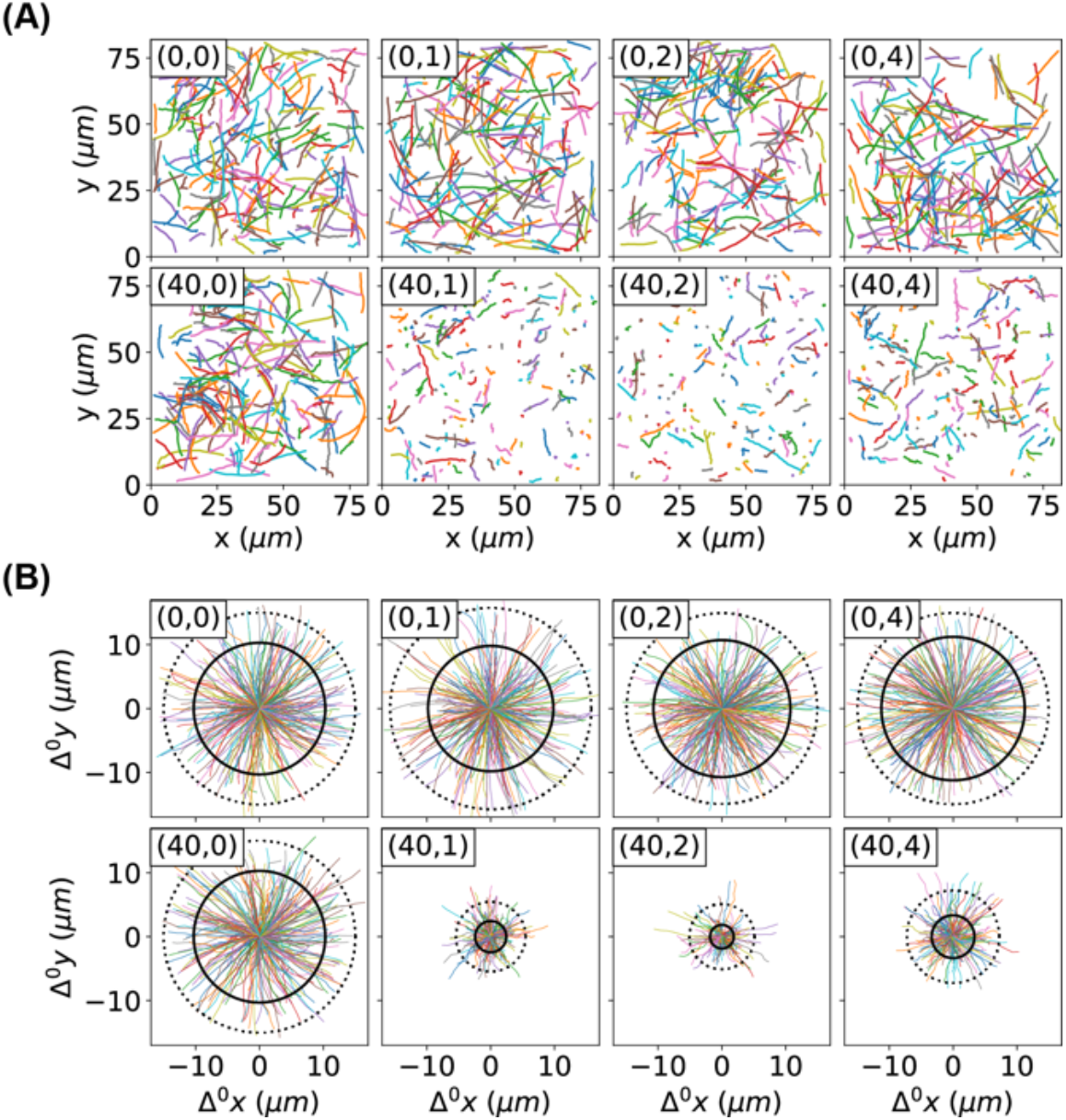
Motion of bacteria. (A) Trajectories of bacteria, untreated or treated by Ag^+^ ions at 30 µM or 40 µM. Each sub-figure contains 200 randomly chosen trajectories, and is labeled by (c_Ag_, T_tr_), where c_Ag_ is the concentration of Ag^+^ ions, and T_tr_ is the treatment/incubation time. (B) Rose graphs of the first 12 frames of trajectories of bacteria, untreated or treated by Ag^+^ ions at 30 µM or 40 µM. Each sub-figure is labeled similar as in panel A. Under each condition, 300 randomly chosen examples of the trajectories were shown in color, while the mean and 90^th^ percentile of the displacements of the first 12 frames of all the trajectories were shown as solid and dotted circles, respectively.

The slower motion of bacteria caused by Ag^+^ ions was further visualized in Fig. 2A by plotting the radii of the mean circles in the rose graphs (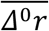, Fig. 1B) as functions of treatment time. To further confirm that the Ag^+^ ions inhibits the movement of bacteria, we calculated the instantaneous velocities of the bacteria directly from the trajectories, *ν* = |**v**| = |*Δ***r**/*Δt*| where *Δt* =0.054 s is the time interval between adjacent frames. The dependence of the mean velocity on the treatment time is shown in Fig. 2B, showing the same trends as 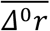. In addition, we examined the distributions of the bacterial velocities (Fig. 2C), and observed a double-peak distribution (centered around 10 and 22 µm/s) for the untreated bacteria (t = 0 hr), while Ag^+^-treatment moved the peak to ~2 µm/s (Fig. 2C). Such significant shift in the velocity-distribution was absent in the negative controls (inset of Fig. 2C). We also found that a second peak/shoulder (7–10 µm/s) emerged in the distributions of bacterial velocities at 4 hr (Fig. 2C), consistent with the previously observed recovery of the bacteria[6, 15].

**Figure 2.**
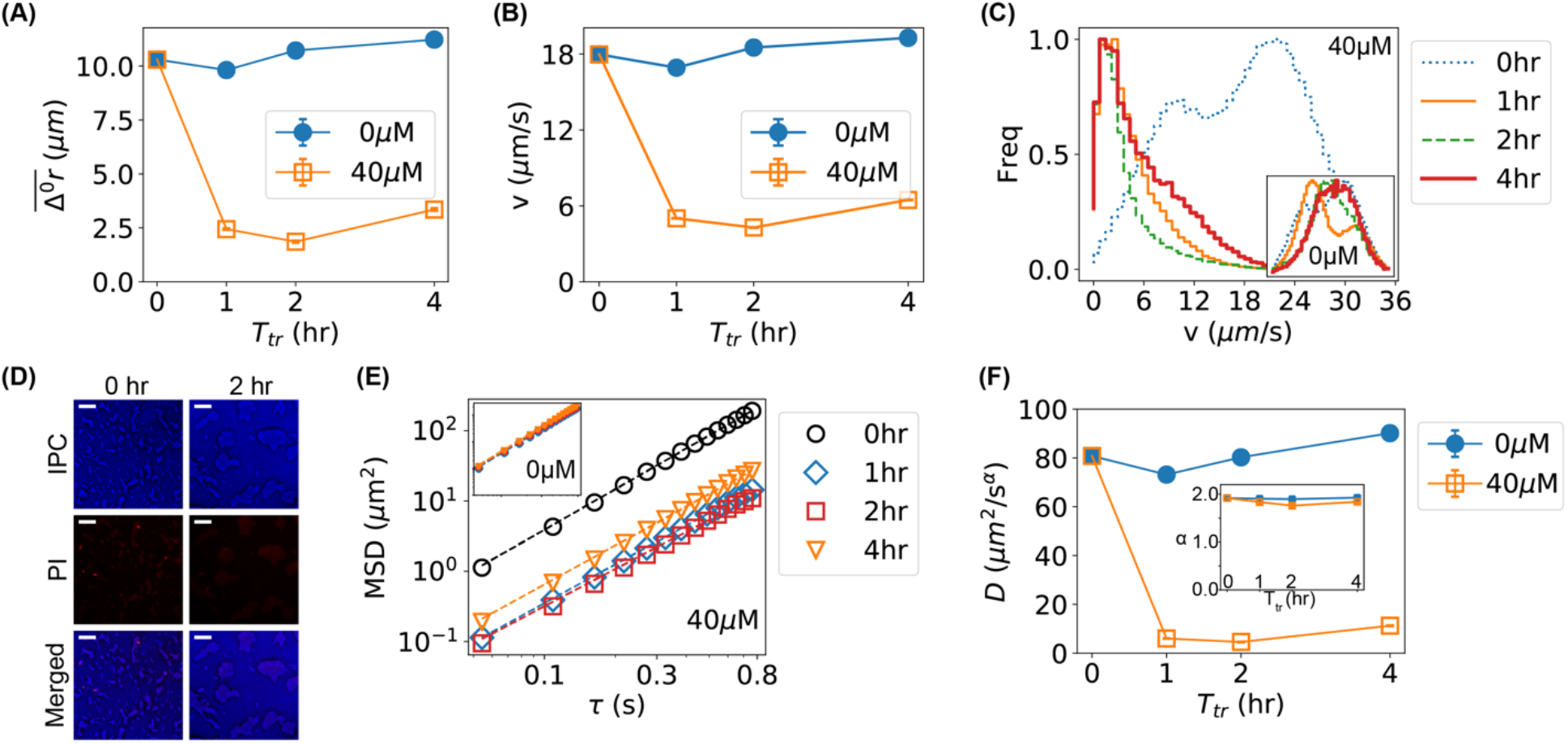
Lower motility of bacteria caused by Ag^+^ ions. (A) The dependence of the mean displacements 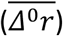 of the first 12 frames of all trajectories of bacteria on incubation/treatment time in the absence (0 µM) and presence of Ag^+^ ions (40 µM). (B) The dependence of the mean bacterial velocity on incubation/treatment time in the absence (0 µM) and presence of Ag^+^ ions (40 µM). (C) Distributions of bacterial velocities in the presence of Ag^+^ ions at 40 µM for 0, 1, 2, and 4 hr. Inset: the corresponding result for untreated bacteria (0 µM). (D) Cell viability assay based on propidium iodide (PI) staining for untreated (0 hr, left column) and treated (2 hr, right column) bacteria. Top: inverted phase-contrast (IPC) images; Middle: fluorescence images due to PI staining; Bottom: merged IPC/PI images. Scale bar = 16 µm. (E) Log-log plot of mean-square-displacements (MSD) *vs.* lag time (*τ*) for trajectories of treated bacteria by Ag^+^ ions at 40 µM for 0, 1, 2, and 4 hr. Inset: the corresponding result for untreated bacteria (0 µM). (F) Dependencies of the generalized diffusion coefficient D and the anomalous scaling exponent *α* (inset) on the incubation/treatment time T_tr_.

As the bacterial velocities after Ag^+^-treatment were close to 0 (peaked at ∼ 2 µm/s), one possibility is that the bacteria were killed by the bacteria at the given concentrations (40 µM) of Ag^+^ ions. However, this possibility was not favored for the following reasons. First, our previous work showed that the majority of bacteria treated with 60 µM Ag^+^ ions were alive, fighting against damages caused by Ag^+^ ions and showing oscillations in their cell-lengths within 12 hr[15]. Second, cell viability assay based on propidium iodide staining[60] showed that the majority of treated bacteria were alive at 40 µM Ag^+^ ions (Fig. 2D). Third, if the bacteria were killed, they would display random diffusion (Brownian motion) and the corresponding mean-square-displacement (MSD) would be proportional to the diffusion coefficient (*D*) and the lag time (*τ*) and shows a slope of 1 in the log-log plot of MSD *vs. τ* (Fig. 2E) [61, 62]; however, fitting the experimental MSD curves (Fig. 2E) with *MSD* = 4*Dτ*^*α*^ (*α* is the anomalous scaling exponent) showed that *α* remained ≈2 in the presence of Ag^+^ ions for various amounts of time (inset of Fig. 2F), indicating that the bacteria retained active motion after Ag^+^-treatment[31, 63]. In contrast, the diffusion coefficient decreased significantly (Fig. 2F), following the same dependence on treatment time as the mean velocity of the bacteria (Figs. 2A and 2B).

### Comparison of bacterial movement before and after Ag^+^-treatment

We quantitatively compared the movement of bacteria before and after Ag^+^-treatment by examining the velocity autocorrelation. Briefly, we calculated the autocorrelations of the x and y components of bacterial velocities, 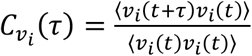, where *ν*_*i*_ = *ν*_*x*_ or *ν*_*y*_ and *τ* is the lag time. For the untreated bacteria, the velocity autocorrelation did not change at different incubation time (insets of Figs. 3A and 3B); in contrast, treating the bacteria with Ag^+^ ions resulted in shifts to the left in the velocity autocorrelation (Figs. 3A and 3B). The left-shift of the velocity autocorrelation suggested that the “persistence” time of the bacterial movement became shorter after Ag^+^-treatment, and the movement of bacteria became not as straight as that before treatment.

**Figure 3.**
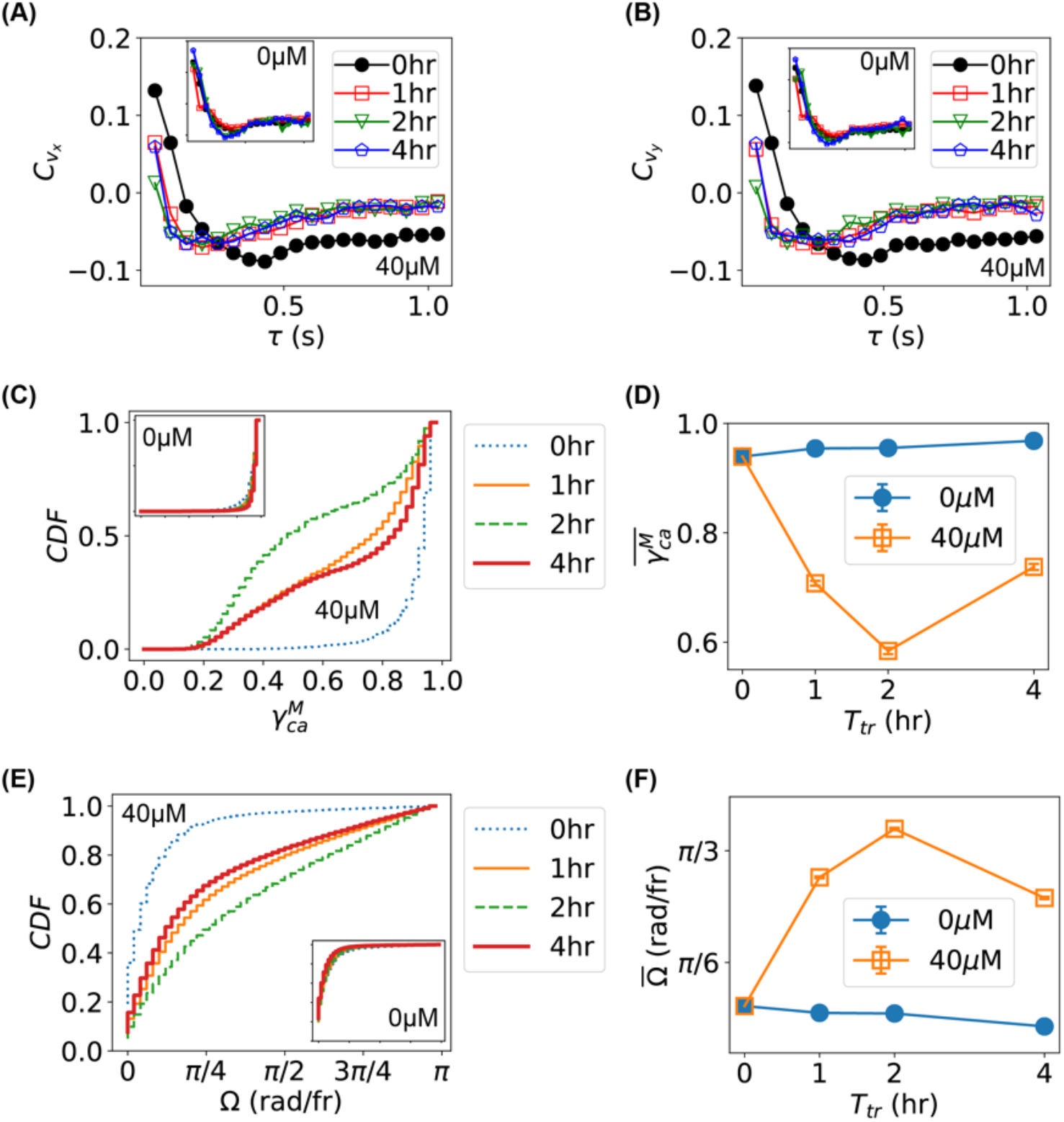
Characterization of bacterial movement and comparison between untreated and treated bacteria. (A, B) Autocorrelation of velocities (A: v_x_; B: v_y_) for bacteria treated with Ag^+^ ions at 40 µM for 0, 1, 2, and 4 hr. Insets: the corresponding results for untreated bacteria. (C) Cumulative distribution function (CDF) of the maximum chord-to-arc ratio 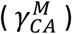 for the trajectories of bacteria untreated (0 hr) or treated with 40 *μ*M Ag^+^ ions for 1, 2, and 4 hr. (D) Dependence of the mean of 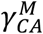 on treatment time. (E) CDF of the changing rate of swimming directions (*Ω*) for bacteria untreated (0 hr) or treated with 40 *μ*M Ag^+^ ions for 1, 2, and 4 hr. (F) Dependence of the mean of *Ω* on treatment time.

We also examined the maximum chord-to-arc ratio 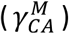 of the trajectories (inspired by TumbleScore[43]), 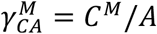, where *C*^*M*^ = max_*i,j*_(|**r**_*i*_ − **r**_*j*_|) is the maximum chord length of a trajectory and *A* = ∑_*i*_|**r**_*i* + 1_ − **r**_*i*_| is the “arc” length of the trajectory. If a trajectory is straight, 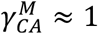, while a trajectory dominated by directional changes gives 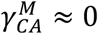; therefore, the maximum chord-to-arc ratio could be used as another indicator of the persistence of the trajectories. The cumulative distributions (CDF) of the 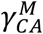 of all the trajectories for bacteria untreated (0 µM and/or 0 hr) or treated with Ag^+^ ions for 1, 2, and 4 hr are shown in Fig. 3C. Compared to the untreated bacteria, the CDFs for treated bacteria rose up at lower 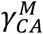 values, indicating that Ag^+^ ions led to higher fractions of lower 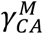. This change was obvious by examining the time dependence of the mean values of 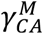 (Fig. 3D). Note that a similar result was observed for the normalized maximum chord-to-arc ratio 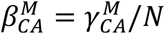 where *N* is the length of the trajectory.

Furthermore, we estimated the changing rate of moving directions directly from the trajectories, *Ω* = cos^−1^(**v**_*i*+1_ ⋅ **v**_*i*_/*ν*_*i*+1_*ν*_*i*_) [43, 44, 45]. The CDFs of *Ω* for all the trajectories of bacteria untreated (0 µM and/or 0 hr) or treated with Ag^+^ ions for 1, 2, and 4 hr are shown in Fig. 3E. We found that the CDFs lowered down after Ag^+^-treatment, indicating increased fraction of higher *Ω* values. This was confirmed by the time dependence of the mean values of *Ω* (Fig. 3F). All the three quantifications (*C*_*ν*_, 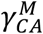, and Ω) showed consistent result that the movement of bacteria became less persistent (i.e., less straight) after subjecting the bacteria to Ag^+^ ions.

### Higher frequency of bacterial tumbling caused by Ag^+^ ions

To further understand the underlying mechanism of the inhibition of bacterial motility by Ag^+^ ions, we exploited the tethering assay on individual bacteria[46, 47]. Briefly, bacteria were tethered to clean glass coverslips through their flagella using biotinylated Anti-FliC antibody, neutravidin, and biotinylated bovine serum albumin (BSA) (Fig. 4A)[48]. The tethered bacteria would rotate on the glass surfaces as the flagellar motors rotate (Movie M3)[48, 64]. Between continuous rotations (i.e., running), occasional pauses and reversed rotations were observed, corresponding to the tumbling of the bacteria (Movie M4)[65, 66]. After monitoring the rotation of the bacteria for 10,000 frames, Ag^+^ ions were added into the samples at a final concentration of 40 µM. The rotation of the bacteria was then monitored for 50,000 to 100,000 frames. It was observed that the rotation of the bacteria slowed down, and that the frequency of pauses increased (Movie M4).

**Figure 4.**
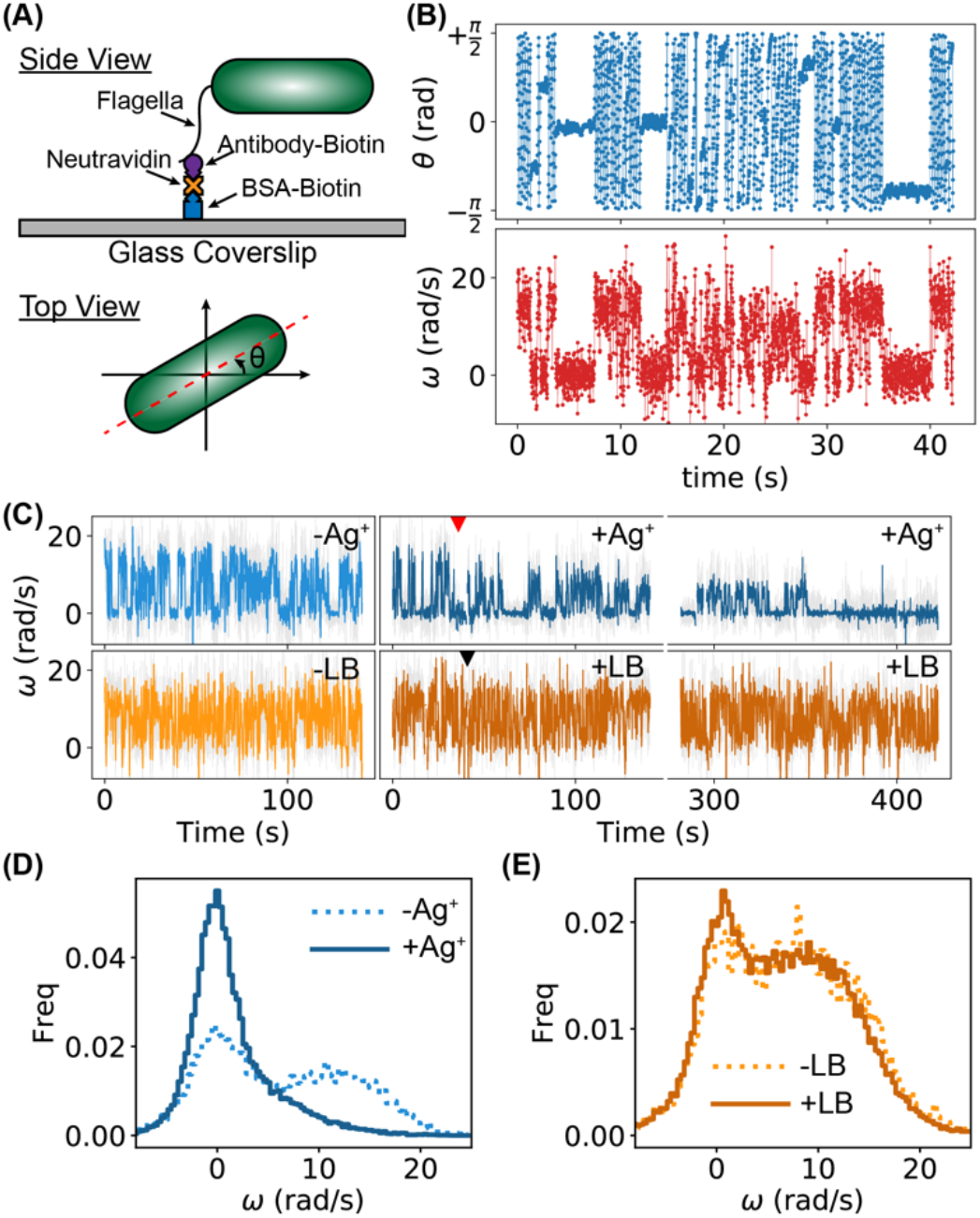
Tethering assay for investigating the running and tumbling of individual bacteria. (A) Tethering of a bacterium on a glass coverslip (side view), and orientation of a bacterium *θ* (top view). (B) Examples of trajectories of orientation *θ* and angular velocity *ω* of a bacterium for 3000 frames (or 42.3 s). (C) Examples of *ω*-trajectories for two bacteria. The top one was treated (blue curves) with Ag^+^ ions; the red arrow indicates the time of adding Ag^+^ ions. The bottom trajectories (orange curves) were for a bacterium without treatment. LB medium was added into the sample at the time indicated by the black arrow. (D) Distributions of *ω* for a bacterium treated by Ag^+^ ions: pre-Ag^+^ (dotted) and post-Ag^+^ (solid). (E) Distributions of *ω* for an untreated bacterium: pre-LB (dotted) and post-LB (solid).

To quantify the results of the tethering assay, we first extracted the orientation of the bacteria, *θ* ∈ (−*π*/2, +*π*/2], in each frame of the movies; then the angular velocities of the bacterial rotations were calculated, *ω* = *Δθ*/*Δt*, where *Δθ* and *Δt* = 0.0141 s were the change of the bacterial orientation and time interval between adjacent frames, respectively. Examples of trajectories of *θ* and *ω* for 3,000 frames (or 42.3 s) for a bacterium before Ag^+^-treatment are shown in Fig. 4B. Two distinct states were observed in the *ω* -trajectory, presumably corresponding to the running and tumbling states[65, 66]. The full *ω*-trajectory (10,000 frames) of Fig. 4B is shown in Fig. 4C (-Ag^+^, light blue), while two segments (each with 10,000 frames) of the *ω*-trajectory of the same bacterium during and after Ag^+^-treatment are also presented (Fig. 4C, +Ag^+^, dark blue), where the red arrow indicates the time of adding Ag^+^ ions. It is clear that the tumbling state (i.e., lower angular velocity) became more frequent after Ag^+^-treatment. In contrast, untreated bacteria (adding LB medium instead of Ag^+^ ions) did not show observable differences in the *ω*-trajectories (Fig. 4C, ±LB, light and dark orange). This observation was quantified by the distribution of the angular velocities. For the control, double peaks were observed for both before and after the addition of LB medium (Fig. 4E); in contrast, the tumbling peak (lower *ω*) became dominant after the addition of Ag^+^ ions (Fig. 4D). The observed increase in the tumbling frequency is consistent with a previous report based on swimming assays for the effect of Ag nanoparticles[28].

### Stalling of flagellar motors caused by Ag^+^ ions

To obtain a deeper understanding of why Ag^+^ ions inhibit the bacterial movement and induce higher tumbling frequency, we performed hidden Markov model (HMM) analysis[55] on the trajectories of angular velocities from the tethering assay. It is noted that hidden Markov model is necessary because the motility states of the bacteria were not directly measured from the experiments; instead, the observable (i.e., the directly measured quantity) was the angular velocity (*ω*). Therefore, our hidden Markov model assumes two states: a running state (R) and a tumbling state (T), which emit observations of angular velocities (Fig. 5A). The probabilities for a bacterium to be in the running and tumbling states are *P*_*R*_ and *P*_*T*_, respectively. The bacterium can switch between the two states, with transition rates of *k*_*RT*_ (from R to T) and *k*_*TR*_ (from T to R). For a given time interval between observations (*Δt* = 0.0141 s between adjacent frames in the tethering assay), the transition probabilities would be *P*_*RT*_ = *k*_*RT*_*Δt* and *P*_*TR*_ = *k*_*TR*_*Δt*, respectively. For each bacterium, we fitted/trained the HMM using the pre-Ag^+^ or pre-LB data, and the fitted model was used to predict the states of all the observed angular velocities for that bacterium, which were then used to estimate the HMM parameters (*P*’s and *k*’s). As an example, the predicted states and the HMM parameters (*P*_*R*_, *P*_*T*_, *k*_*RT*_, and *k*_*TR*_) for the ±Ag^+^ bacterium in Fig. 4C are presented in Figs. 5C and 5B, respectively. Two significant changes were observed. First, the tumbling probability (*P*_*T*_) increased dramatically from 49% to 87% (correspondingly, *P*_*R*_ = 1 − *P*_*T*_ decreased); second, while the running-to-tumbling transition rate increased slightly, the tumbling-to-running transition rate *k*_*TR*_ decreased significantly by > 4-fold after Ag^+^-treatment (3.12 *s*^−1^ to 0.75 *s*^−1^). These observations suggest that Ag^+^ ions lead to higher tumbling frequency by blocking the transition from the tumbling state to the running state.

**Figure 5.**
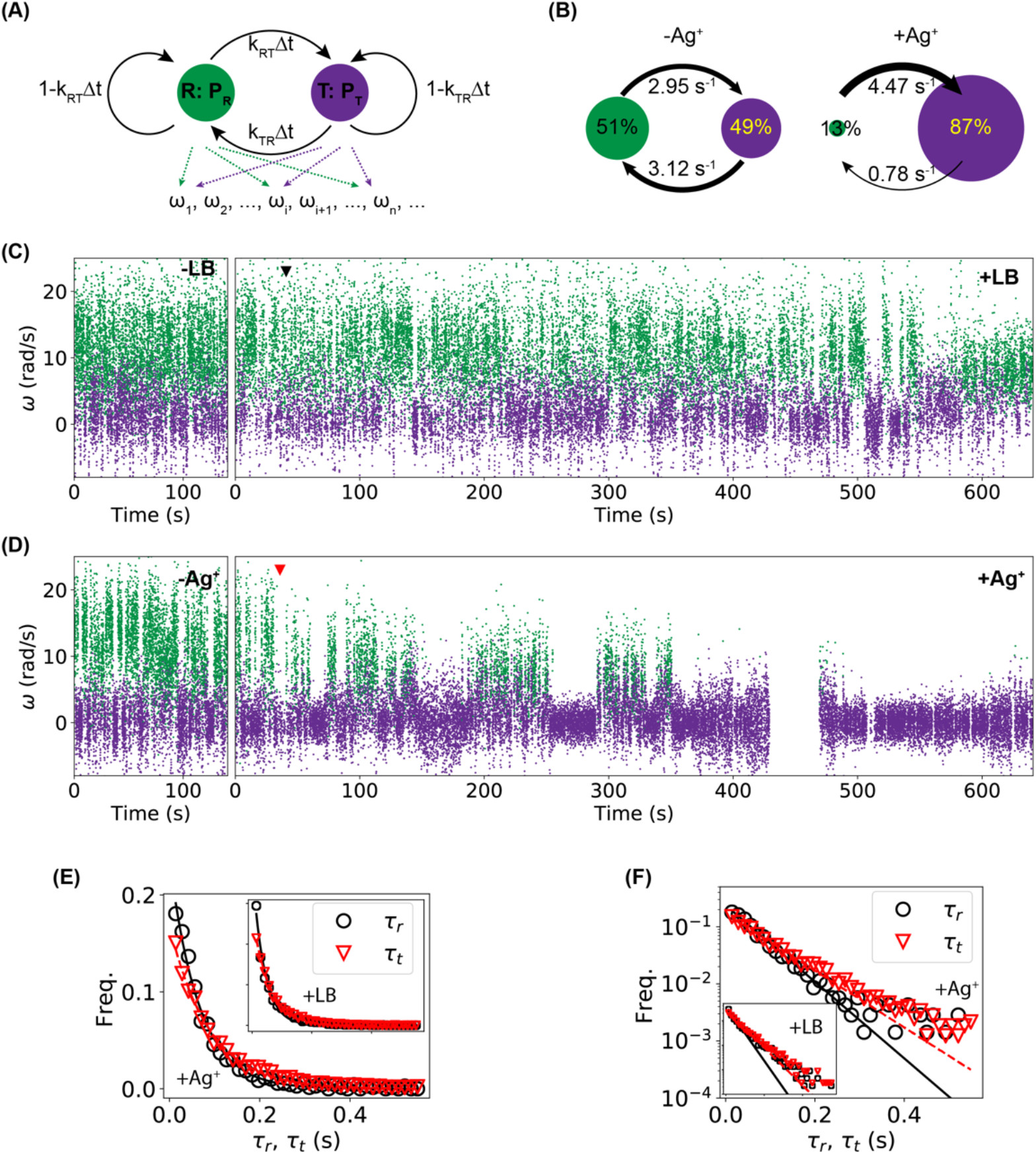
Hidden Markov model (HMM) analysis. (A) The hidden Markov model with two states (running (R) *vs.* tumbling (T)), which emit observations of angular velocities *ω*_*i*_. The probabilities for the system to be in the running and tumbling states are *P*_*R*_ and *P*_*T*_, respectively. The transition probabilities between the two states are *P*_*RT*_ = *k*_*RT*_*Δt* and *P*_*TR*_ = *k*_*TR*_*Δt*, where *k*_*RT*_ and *k*_*TR*_ are the corresponding transition rates and *Δt* is the time interval between observations. (B) Predicted parameters (*P*_*R*_, *P*_*T*_, *k*_*RT*_ and *k*_*TR*_) from the HMM analysis for pre-Ag^+^ and post-Ag^+^ *ω*-trajectories of the bacterium in the top row of Fig. 4C. (C, D) Predictions of states from the fitted/trained HMM model for the angular velocity (*ω*) trajectories for (C) an untreated bacterium and (D) an Ag^+^-treated bacterium. Green and purple dots correspond to the running and tumbling states, respectively. (E, F) Distributions of the dwell times (*τ*_*r*_ for running dwell time and *τ*_*t*_ for tumbling dwell time) from the (E) untreated and (F) Ag-treated bacteria shown in panels (C) and (D). Solid and dashed lines are fitted exponential curves. Insets: the same data plotted in log-linear scale.

As simple and hidden Markov models typically assume exponential distributions for the dwell times (i.e., the time staying in the states), we wondered whether and how this assumption was satisfied in the tethering assay. Briefly, from the predicted states for the control and sample shown in Fig. 5C (+LB and +Ag^+^, respectively), we calculated the running time (*τ*_y_) and tumbling time (*τ*_(_) and found that the distributions of both dwell times followed roughly the exponential distribution for both the control (+LB) and the sample (+Ag^+^), as shown in Fig. 5D, where the solid and dashed lines are fittings. This observation indicates that the hidden Markov model is reasonably suitable for the analysis here. On the other hand, we note that, a closer look on the distributions of the dwell times in the log-linear scale indicated that a single exponential decay did not fit the data well (Fig. 5E), suggesting that modified hidden Markov models that assume arbitrary distributions of the dwell times may improve the analysis.

We replicated the tethering assay experiments and HMM analysis on 10 untreated (±LB) and 15 treated (±Ag^+^) bacteria. We observed large variations in the absolute values of the angular velocities for different bacteria, which could be attributed to differences in the cell length, the number of tethered flagella per bacterium, and the location of tethering points on the flagella[48, 67]. To compare among different bacteria, we used the relative changes in the HMM parameters, 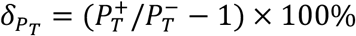 and 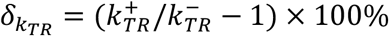, where the superscripts (+ and −) stand for after and before the addition of Ag^+^ ions (or the addition of LB medium for the controls), respectively. The relative changes for the untreated (orange squares) and Ag^+^-treated bacteria (blue circles) from the full-length trajectories are shown in Fig. 6A. Performing one-sample *t*-test showed that the increase in *P*_*T*_ and decrease in *k*_*TR*_ were much more statistically significant for the Ag^+^-treated bacteria (*p*-values: 2.1 × 10^−4^ and 4.1 × 10^−10^ for *P*_*T*_ and *k*_*TR*_, respectively) than the untreated cells (*p*-values: 0.065 and 0.014, respectively). Two-sample *t*-test showed that the differences between the treated and untreated samples were also statistically significant (e.g., the *p*-value for *k*_*TR*_ was 4.2 × 10^−4^).

**Figure 6.**
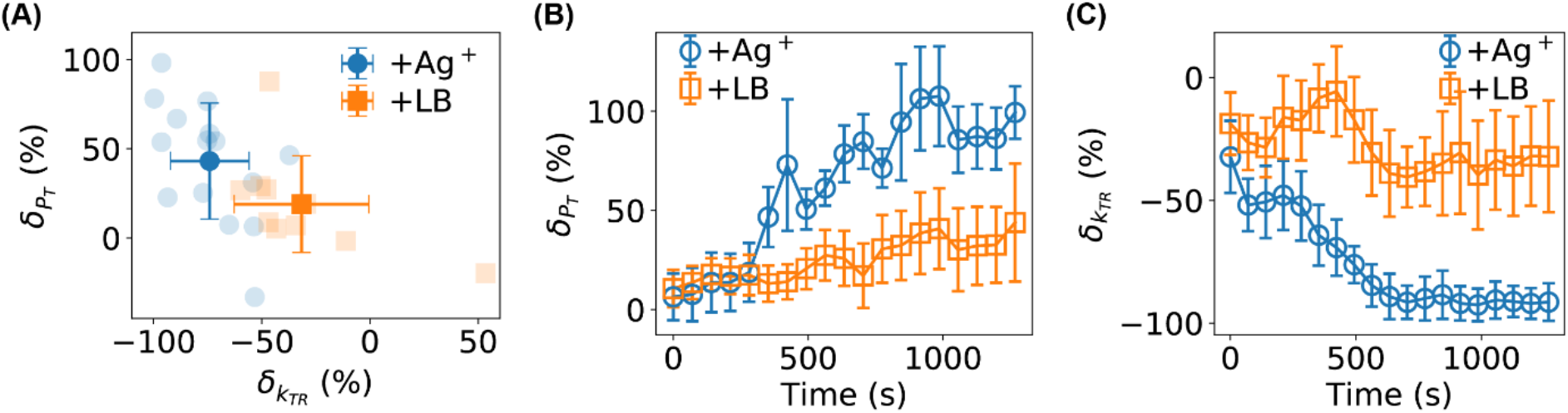
(A) Statistics of the relative changes in *P*_*T*_ and *k*_*TR*_ for 10 untreated (orange squares) and 15 Ag^+^-treated bacteria (blue circles). Error bars stand for standard deviation. (B, C) Time dependencies of the relative changes in (B) *P*_*T*_, and (C) *k*_*TR*_ for untreated (orange squares) and Ag^+^-treated (blue circles) bacteria. Error bars stand for the standard error of the mean.

Finally, we examined the dependence of the HMM parameters on the treatment time (Figs. 6B and 6C), which was done by analyzing individual segments of the full-length *ω* -trajectories (window-size = 10,000 frames, stride between segments = 5,000 frames) using the fitted/trained HMM models. We observed that both 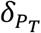 and 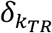 started from ≈ 0, which is reasonable as it takes time for the Ag^+^ ions to diffuse to the bacteria and affect the bacteria. More interestingly, the effects of Ag^+^ ions became more and more significant after ∼300 s compared to the controls (Figs. 6B and 6C). After ~750 s, the relative change of *k*_*TR*_ reached ∼ −90%, suggesting that Ag^+^ ions prevented the flagellar motor of the bacteria from rotating effectively and efficiently.

## Conclusions and Discussions

To conclude, we directly visualized and investigated the antibiotic effects of Ag^+^ ions on the motility of *E. coli* bacteria based on swimming and tethering assays. From the swimming assay, we observed that the bacteria slowed down dramatically when subjected to Ag^+^ ions. Characterization of the swimming trajectories showed higher changing rates of swimming directions. In addition, we tethered the bacteria on glass surfaces through bacterial flagella (i.e., the tethering assay) and monitored the rotation of flagellar motors directly, from which we observed an increase in the tumbling frequency due to Ag^+^-treatment. We performed hidden Markov model (HMM) analysis on the trajectories of angular velocities of the bacterial rotation and compared the bacteria before and after Ag^+^-treatment. It was found that treated bacteria stayed in the tumbling state with much higher probability and that the transition rate from the tumbling state to the running state decreased in the presence of Ag^+^ ions, suggesting that Ag^+^ ions stalled the flagellar motors and prevented them from rotating.

The observed inhibition of bacterial movement and higher frequency of tumbling caused by Ag^+^ ions confirmed our hypothesis that the motility of bacteria is significantly affected by Ag. This work provides direct visualization of the Ag’s effects on the bacterial movements and advances quantitatively our understanding on the mechanism of Ag-based antimicrobial agents in terms of bacterial motility. More importantly, it raises more interesting questions worth further investigations. For example, what is the molecular basis for the observed slower swimming, more frequent tumbling, and motor stalling, when subjecting bacteria to Ag^+^ ions? To what extent the observed effects on the bacterial motility are Ag-specific? How will the bacteria adapt to, or become resistant against, the Ag-induced damages on the bacterial movement? How will bacterial death be related to the observed lower motility? Addressing these biological questions experimentally is expected to be of great importance and interest for understanding the fundamental antimicrobial mechanism of Ag and further exploring their potential biomedical applications.

Our data suggest that the observed effects of Ag^+^ ions on the bacterial motility are likely due to direct interactions between the bacterial flagella and Ag^+^ ions, which can be seen from the response time of the rotation of bacteria to the addition of Ag^+^ ions (Fig. 6) in the tethering assays. In our tethering experiments, as the Ag^+^ ions were added to the top surface of the liquid medium in the Petri-dish above the bacteria under observation, the distance that Ag^+^ ions need to travel to the bacteria is roughly Δ*x* = 0.2 cm (estimated from the volume of the culture medium, 2 mL, and the diameter of the Petri-dish, 3.5 cm). Considering that the diffusion coefficient of Ag^+^ ions in water[68] is in the order of *D* = 1.5 × 10^−5^ cm^2^/s, the time scale for the Ag^+^ ions to reach the bacteria is in the order of 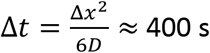, which is close to the response time (300 – 750 s) of bacteria to the Ag^+^ ions that we measured from our tethering assays (Fig. 6B and 6C). If the observed effects of Ag^+^ ions on the bacterial motility were due to indirect interactions, such as those through regulatory proteins and membrane damages, the response time is expected to be longer as time is needed to transduce those indirect effects to the flagellar motor. Therefore, it is suggested to focus on the bacterial flagella when searching for molecular basis for the Ag-caused slower swimming, more frequent tumbling, and motor stalling in future studies.

## Acknowledgement

This work was supported by the University of Arkansas, the Arkansas Biosciences Institute, and the National Science Foundation (Grant No. 1826642 to YW and JC). AR and MK were partially supported by NSF-REU grants (Grant No. 1460754 and Grant No. 1851919).

